# Dynamics in the *Phytophthora capsici* effector AVR3a11 confirm the core WY domain fold

**DOI:** 10.1101/2024.09.04.611235

**Authors:** James Tolchard, Vicki S. Chambers, Laurence S. Boutemy, Mark J. Banfield, Tharin M. A. Blumenschein

**Affiliations:** School of Chemistry, University of East Anglia, Norwich Research Park, Norwich, NR4 7TJ, UK; Department of Biochemistry and Metabolism, John Innes Centre, Norwich Research Park, Norwich, NR4 7UH, UK; Centre de RMN à Très Hauts Champs de Lyon (CRMN), Claude Bernard University Lyon 1, Villeurbanne, 69100, France; Illumina, Illumina Centre, Cambridge, CB216DF, UK; Norwich School, Norwich, NR1 4DD, UK

**Keywords:** effector protein, hydrogen-deuterium exchange, NMR, dynamics, oomycete

## Abstract

Oomycete pathogens cause large economic losses in agriculture through diseases such as late blight (*Phytophthora infestans*), and stem and root rot of soybean (*Phytophthora sojae*). The effector protein AVR3a, from *P. infestans*, and its homologue AVR3a11 from *P. capsici*, are examples of host-translocated effectors that interact with plant proteins to evade defence mechanisms and enable infection. Both proteins belong to the family of RXLR effectors and contain an N-terminal secretion signal, an RXLR motif for translocation into the host cell, and a C-terminal effector domain. Within this family, a large number of proteins have been predicted to contain one or more WY domains as their effector domain, and this domain is proposed to encompass a conserved minimal core fold containing three helices, further stabilised by additional helices or dimerization. In AVR3a11, a helical N-terminal extension to the core fold forms a four-helix bundle, as determined by X-ray crystallography. For a complete picture of the dynamics of AVR3a11, we have determined the solution structure of AVR3a11, and studied its dynamics in the fast timescale (ns-ps, from NMR relaxation parameters) and in the slow timescale (seconds to minutes, from hydrogen/deuterium exchange experiments). Hydrogen/deuterium exchange showed that the N-terminal helix is less stable than the other three helices, confirming the core fold originally proposed. Relaxation measurements confirm that AVR3a11 undergoes extensive conformational exchange, despite the uniform presence of fast motions in the spectral density function throughout most of its sequence. As functional residues are located in the more mobile regions, this flexibility in the slow/intermediate timescale may be functionally important.

**Lay Summary:** The effector protein AVR3a11, from the plant pathogen *Phytophthora capsici*, shows conformational flexibility in solution, particularly in the N-terminal helix, and in an intermediate timescale (ms-s). This confirms the core fold originally proposed and suggests that flexibility may be functionally important.

## Introduction

Filamentous pathogens of plants, including fungi and oomycetes, are responsible for large economic losses in agriculture. The oomycete *Phytophthora infestans*, the causative agent of late blight, is responsible for yield losses worth over € 10 billion a year worldwide (1). Meanwhile, losses caused by *P. capsici*, a pathogen of peppers, tomato, eggplant and cucurbits (such as squashes and cucumber), have increased significantly in the last few decades (2).

*Phytophthora* pathogens obtain nutrients from host plant cells by forming haustoria, specialised structures that penetrate the plant cell wall without rupturing the cell membrane (3). Specific molecular interactions between plant and pathogen are essential not only for successful infection, but also for plant resistance against disease. The first layer of plant defence against pathogens is initiated at the cell surface by pattern recognition receptors (PRRs), which detect the presence of pathogen-associated molecular patterns (PAMPs). Adapted pathogens of plants produce effector proteins, some of which interfere with PRR-mediated immunity (4, 5). These effectors, or their activities, can be detected by intracellular nucleotide binding-leucine rich repeat (NLR) receptors, triggering a second layer of plant defence. Effectors that trigger NLR-mediated immunity are often termed avirulence (AVR) proteins and their recognition can result in the hypersensitive response (HR) and programmed cell death (PCD) (6, 7).

The well-studied *P. infestans* effector AVR3a activates gene-for-gene HR in plants expressing the NLR protein *R3a* (8). Close homologues of AVR3a are found in a number of *Phytophthora* species, including *P. capsici* and *P. sojae* (pathogen of soybean) (9). AVR3a suppresses programmed cell death induced by the *P. infestans* elicitin INF-1, and other PAMPs, by at least two mechanisms: stabilising the plant U-box E3 ubiquitin ligase CMPG1 (involved in cell death triggered by a number of PAMPs) (10-12), and associating with a plant GTPase dynamin-related protein 2 (DRP2), a vesicle trafficking protein involved in receptor-mediated endocytosis (13). AVR3a and its homologues are composed of three distinct regions: an N-terminal signal region, which determines secretion from *P. infestans*; a predicted disordered RXLR motif, with a role in effector delivery (14); and a C-terminal domain, responsible for effector activity (15). AVR3a occurs naturally as two predominant alleles, AVR3a^KI^ and AVR3a^EM^, differing in two amino acid positions: 80 (Lys or Glu) and 103 (Ile or Met) (8). AVR3a^KI^ (Lys80/Ile103) is recognised by *R3a* in potato and the model solanaceous plant *N. benthamiana*, leading to HR, whilst AVR3a^EM^ (Glu80/Met103) evades recognition (8). Point mutations in R3a (termed R3a^+^) can enable recognition of AVR3a^EM^ in *N. benthamiana* (16). The regions of AVR3a responsible for recognition by R3a and interaction with CMPG1 are at least partially independent. The C-terminal residue, Tyr147, is essential for CMPG1 stabilisation and suppression of CMPG1-dependent cell death, but is not required for recognition by R3a (11, 17). However, whilst AVR3a^EM^ escapes recognition by R3a, it does not stabilise CMPG1 or suppress INF1-mediated cell death to the same level as AVR3a^KI^, suggesting an overlap between the regions involved in the two functions (17). Other Avr3a-like effectors have been shown to target the plant CAD7 subfamily, cinnamaldehyde dehydrogenases which act as plant immunity regulators (18).

A high-resolution crystal structure of the effector domain of *P. capsici* AVR3a11 (residues Thr70 to Val132) was the first crystal structure of an effector in this family to be reported (9), revealing a four-helical bundle fold. Structural homology to the dimeric *P. infestans* RXLR effector PexRD2 and sequence motif analysis in three other *Phytophthora* species revealed a three-helix folding unit, the “WY” domain, named after the conserved Trp and Tyr residues in its hydrophobic core. This domain leads to a number of possible structural arrangements, from a homodimer in which the WY core interactions happen across the dimer interface (PiSFI3) (19) to a series of WY repeats which cross-stabilise each other (PexRD54, PsAvh240) (20, 21), and even to an extended LWY fold, where an extended version of the oomycete L-motif connects and stabilises the WY motif (22). Taken together, the variety of structures reinforces the versatility of the WY domain as a basis for effector evolution (23). Almost half of *Phytophthora* RXLR effectors, and a quarter of RXLR effectors in the Arabidopsis pathogen *Hyaloperonospora arabidopsidis*, are predicted to contain WY domains (9).

In AVR3a11, the archetypal core three-helix fold is further stabilised by a helical N-terminal extension, forming the four-helical bundle. This protein fold is conserved between different oomycete effectors, despite a lack of sequence similarity (24). Solution structures of AVR3a (14) and AVR3a4 (25) revealed very similar structures to AVR3a11 for these close homologues.

Preliminary nuclear magnetic resonance (NMR) experiments of the predicted effector domain of AVR3a11 (residues Gly63 to Val132) showed extensive conformational exchange in this longer construct (9). This suggests that dynamics are a key property of this protein, and may be important for function, not only of AVR3a11 but also of homologous effectors. Deletion of 7 extensively broadened residues at the N-terminus led to the shorter construct, AVR3a11_70-132_, used for determination of the crystal structure (9). The other region in conformational exchange, the loop between helices 3 and 4, is stabilised by crystal contacts in the published structure. To compare the structural features in solution, and to investigate the dynamics of AVR3a11, here we present the solution structure for AVR3a11_63-132_, determined by NMR spectroscopy, and dynamics measurements for the shorter construct AVR3a11_70-132_. The main structural features are all preserved, although helix 4 is a couple of residues longer, towards the C-terminus. The dynamics of AVR3a11_70-132_ were studied both in the ps-ns timescale, calculated from the measurement of NMR relaxation parameters, and in a slower timescale (seconds to minutes), measured from hydrogen/deuterium exchange experiments. While low-amplitude fast movements are uniform throughout the helical bundle, different helices have different stabilities in the slower timescale. Helix 1 in particular, the N-terminal extension to the WY domain, is less rigid than the core three helices. Some characteristics of the solution structure and dynamics of AVR3a11 are comparable to other WY domain effectors studied by NMR, and are likely a general property of these proteins.

## Materials and methods

### Sample preparation

Previously described AVR3a11 constructs (9) were used for protein expression. ^15^N-labelled AVR3a11_70-132_ (residues Thr70-Val132) was expressed in *E. coli* BL21(DE3) grown in N-5052 autoinduction media (26) overnight at 30 °C. [^13^C, ^15^N]-labelled AVR3a11_63-132_ (residues Gly63-Val132) was expressed in *E. coli* BL21*(DE3) grown in M9 minimal medium supplemented with trace elements, MEM vitamin solution (Sigma-Aldrich), 0.2% ^13^C-glucose and 20 mM ^15^NH_4_Cl, for 3 hours of post-induction growth at 37 °C (cells induced at A_600_ ∼ 0.4 – 0.6 with 1 mM IPTG). ^15^N- and [^13^C, ^15^N]-labelled AVR3a11 constructs were purified as described (9), in the presence of Complete EDTA-free protease inhibitor cocktail (Roche Diagnostics). In both cases, the His-tag was removed by cleavage with 3C protease, leaving two residues from the linker sequence in the amino terminus of the constructs (Gly-Pro). Purified AVR3a11 was concentrated to approximately 1 mM in 90% H_2_O/10% D_2_O, 10 mM sodium phosphate pH 8.8, 50 mM sodium sulphate, 0.03% sodium azide, 0.2 mM 2,2-dimethyl-2-silapentane-5-sulfonic acid (DSS), and Complete EDTA-free protease inhibitor cocktail (Roche Diagnostics) in the recommended concentration.

### NMR spectroscopy

The following spectra were acquired in a Bruker Avance III 800 MHz NMR spectrometer with a TXI probe with Z-pulsed field gradients at 298 K for backbone and side chain assignment, and measurement of distance restraints for structure calculation, using [^13^C, ^15^N]-labelled AVR3a11_63-132_: ^15^N-HSQC, ^13^C-HSQC, ^13^C-TROSY-HSQC in the aromatic region (27), CBCA(CO)NH, CBCANH (28, 29), CC(CO)NH, H(CCO)NH (30), ^15^N-TOCSY-HSQC, ^15^N-NOESY-HSQC (mixing time of 100 ms), ^13^C-NOESY-HSQC (mixing time of 120 ms) (31) and (H)CB(CGCC-TOCSY)H^ar^ (32). Backbone amide relaxation experiments – ^15^N T_1_, ^15^N T_2_, and [^1^H]^15^N NOE (33) – were acquired at 293 K in a Bruker Avance III 800 MHz and a Bruker Avance I 500 MHz NMR spectrometers using ^15^N-labelled AVR3a11_70-132_. T_1_ measurements were performed with a recovery delay of 4 s, and relaxation delays of 0.02, 0.1, 0.2, 0.5, 0.75, 1, and 4 s. The relaxation delay of 1 s was repeated to evaluate data consistency. T_2_ measurements were performed with a recovery delay of 4 s at 500 MHz, and 5 s at 800 MHz, and relaxation delays of 17, 51, 85, 136, 170, 204 and 254 ms. Relaxation delays of 17, 85 and 170 ms were repeated to evaluate data consistency. NOE measurements used a saturation delay of 4 s, replaced by a relaxation delay of 4 s in the reference experiment. All NMR spectra were processed using NMRPipe (34) and analysed with the CcpNmr Analysis package (35). ^1^H referencing for all NMR spectra was performed using the internal DSS reference. ^15^N and ^13^C were referenced according to the ratio of their gyromagnetic ratios to ^1^H, as described (36).

### Structure calculation

Backbone and side chain resonance assignments were obtained manually from the NMR spectra, and converted to XEASY (37) format with CcpNmr FormatConverter (35). Backbone assignments were used to generate dihedral angle restraints for 49 residues with the TALOS+ webserver (38). ^15^N-NOESY-HSQC and ^13^C-NOESY-HSQC spectra were converted to CARA (39) format, and used as input to the UNIO (40) software package together with the dihedral angle restraints. UNIO uses the ATNOS/CANDID algorithms (41, 42) to pick and assign NOE crosspeaks. Using peak picking tolerances of 0.03 ppm (^1^H) and 0.4 ppm (^13^C, ^15^N), and other default parameters, seven iterative cycles of NOE crosspeak assignment, restraint refinement and structure calculation were used within UNIO to obtain a final list of 1020 distance restraints, with the molecular dynamics algorithm CYANA 2.1 (43). These distance restraints, combined with the dihedral angle restraints, were used for the calculation of 100 structures followed by water minimisation within CNS 1.3 (44, 45), using the RECOORD protocol (46). The 20 structures with lowest energy and no violations greater than 0.5 Å for NOE or 5° for dihedral angles were selected to form the final structural ensemble, which was validated in PSVS (Protein Structure Validation Software Suite (47)), including the validation tools MolProbity (48) and PROCHECK (49).

### Hydrogen/deuterium exchange

Hydrogen/deuterium exchange was performed using a ^15^N-labelled AVR3a11_70-132_ sample prepared as described above. Imidazole at 1 mM was added to the sample to monitor the pH (50). The sample’s pH was adjusted to 6.87 with HCl, then the sample was lyophilised and resuspended in ice-cold deuterium oxide. Loss of signals was followed with [^1^H, ^15^N]-SOFAST-HMQC (51) spectra recorded at 278 K in a Bruker Avance III 800 MHz spectrometer. Spectra were recorded at 4, 7, 10, 13, 16, 19, 22, 25, 30, 36, 46, 57, 71, 86, 101, 116, 146, 176, 206, 236, 296, 356, 416, 536, 656, 776, 956, 1196, and 1440 minutes after the addition of deuterium oxide. Single exponential decay curves were fitted to the peak intensities, adjusted for the number of scans in the spectrum and with uncertainties estimated from the standard deviation of the noise in a blank spectral region in nmrDraw (34), using the first order exponential decay fit in the software package Origin (OriginLab, Northampton, MA, USA). Protection factors were calculated using random coil exchange rates and temperature correction as described by Bai *et al*. (52).

### Relaxation analysis

Peak lists for every relaxation experiment were exported from CcpNmr Analysis in NMRView (53) format. The standard deviation of the spectral noise was estimated in a blank region of each spectrum in nmrDraw (34). Peak intensities and noise estimates were used in Relax (54) to fit T_1_ and T_2_ to a single exponential decay function, and to calculate the heteronuclear NOE ratio. The experimental backbone amide relaxation parameters were fitted according to the Lipari-Szabo approach (55, 56), using five different models for the spectral density function (S^2^-τ_m_, S^2^-τ_m_-τ_e_, S^2^-τ_m_-R_ex_, S^2^-τ_m_-τ_e_-R_ex_, and a two-time scale model). Fitting was performed with ModelFree4 (57) and FastModelFree (58), considering Avr3a11_70-132_ axially symmetric, and using a diffusion tensor calculated from the crystal structure (9) and the relaxation data with the programme Quadric_Diffusion (59). Reduced spectral density mapping (60) was calculated using Relax (54) with default parameters.

## Results

### Structure determination

Two constructs for the effector domain of *P. capsici* AVR3a11, AVR3a11_63-132_ (containing an extra N-terminal 7 residues, predicted to be helical) and AVR3a11_70-132_ (corresponding to the crystal structure), were expressed in *E. coli* and purified for NMR studies. Both AVR3a11 constructs produced well-dispersed [^1^H, ^15^N]-HSQC NMR spectra, as shown in Fig. 1. After accounting for overlapped crosspeaks, 15 residues in AVR3a11_63-132_ could not be observed, corresponding mostly to the N-terminal extension in relation to the crystal structure, and to loop 3 (between helices 3 and 4). Overall, 77% of the amides, 69% of the remaining backbone nuclei and 71% of the non-labile protons were visible and assigned in AVR3a11_63-132_ (Fig. S1 in Supplementary Material). The assignments were deposited to the Biological Magnetic Resonance Data Bank (BMRB) (61) under accession code 18910. Dihedral angle restraints for phi and psi angles were obtained from ^1^H, ^13^C and ^15^N chemical shifts, and distance restraints for structural calculation were determined from NOESY crosspeaks. An ensemble of 20 refined structural models was calculated, obtaining good validation scores (Table S1), and was deposited in the Protein Data Bank (PDB) under accession code 3ZGK. The final structural models (Fig. 2) adopt a four-helix bundle conformation, with two pairs of anti-parallel α-helices. RMSD for the ordered regions are 0.75 ± 0.12 Å for backbone atoms, and 1.35 ± 0.14 Å for heavy atoms.

**Figure 1.**
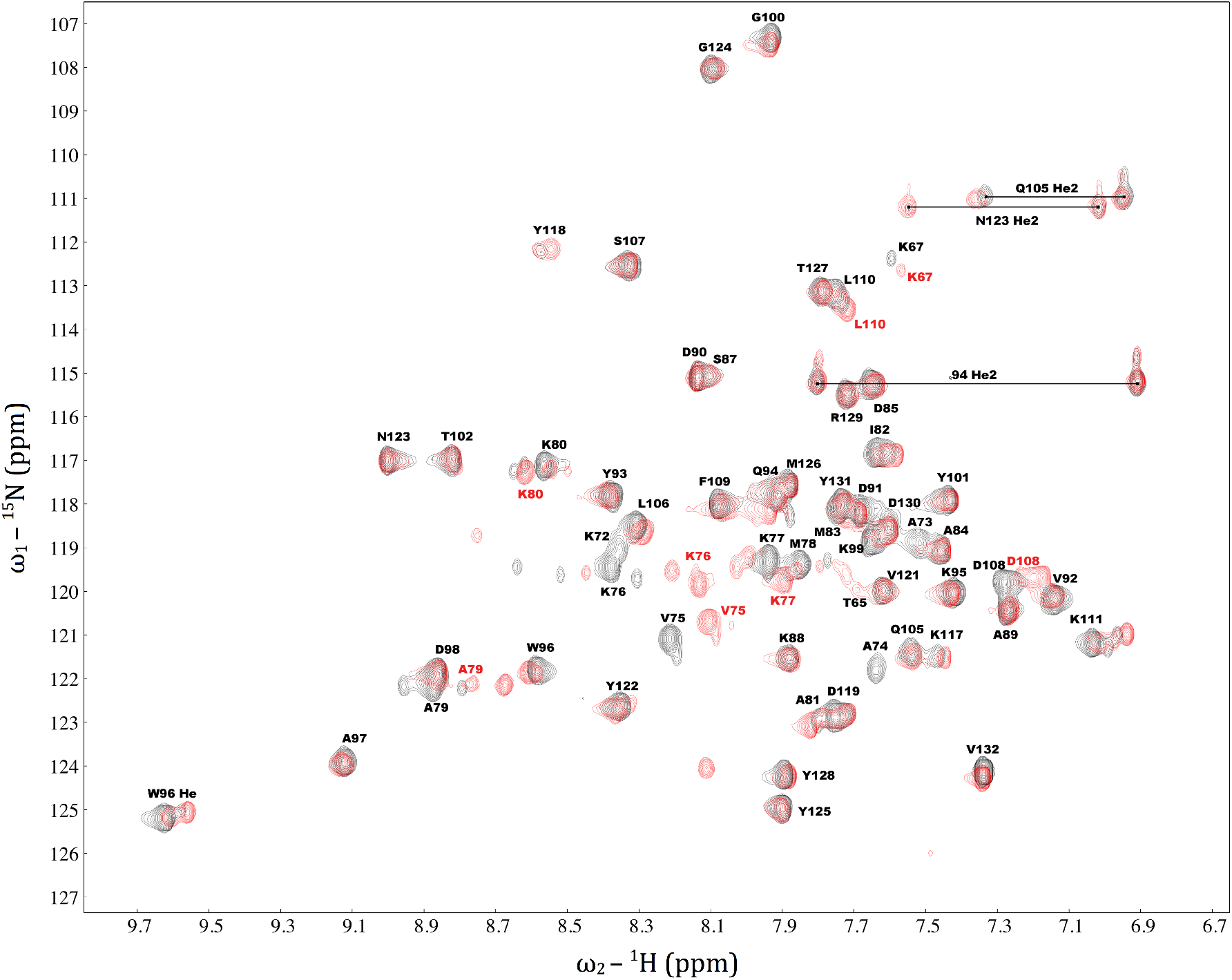
Two-dimensional [^1^H, ^15^N]-HSQC NMR spectra of AVR3a11_63-132_ (black) and AVR3a11_70-132_ (red). Minor peaks corresponding to alternative conformations of AVR3a11 can be observed, for instance, in the proximity to Lys76, Ala79 and Lys111.

**Figure 2.**
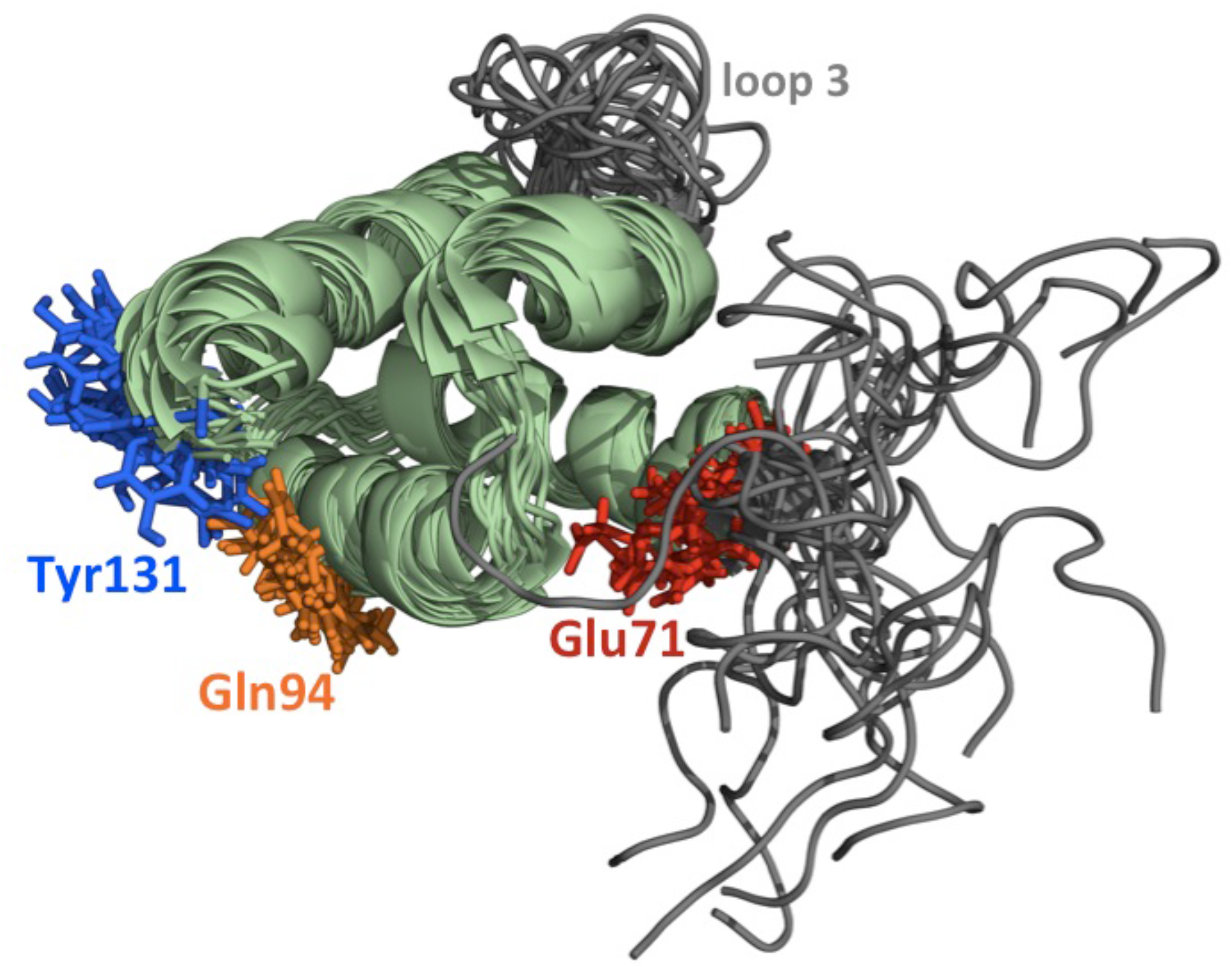
Ensemble of solution structures for AVR3a1163-132, showing the 20 lowest-energy structures calculated. The side chains of equivalent positions to functionally relevant residues in the homologue *P. infestans* AVR3a are highlighted: Glu71 and Gln94 (recognition by the host cell resistance protein R3a) and Tyr131 (interaction with CMPG1 and inhibition of programmed cell death).

Due to an absence of observable peaks, no structural restraints were defined for residues 63-69 or loop 3, leading to significant heterogeneity in these regions across the structural ensemble. The inability to observe these regions is likely due to conformational exchange in the intermediate regime on the chemical shift timescale, and indicates that neither of these regions adopt a single, well-folded conformation in solution. Conformational exchange was also responsible for a few other residues in the protein displaying minor peaks (Fig. S2).

The extensive conformational exchange for residues 63-69, as observed from the NMR spectra, provide a likely explanation for why it was not possible to crystallise AVR3a11_63-132_ (9).

Comparison between the solution structural ensemble of AVR3a11_63-132_ described here and the 0.9 Å resolution crystal structure of AVR3a11_70-132_ (9) (PDB access code 3ZR8) shows good agreement between them, with 1.05 Å RMSD between C_α_s in ordered regions of the most representative conformer and the crystal structure (Fig. 3). This confirms that the shorter construct does not affect the overall structure in AVR3a11_70-132_. The main differences between the solution and crystal structures are located in loop 3, and in the final C-terminal residues. In the solution ensemble, loop 3 displayed a high degree of conformational heterogeneity across the individual models. Whilst this is a consequence of unrestrained molecular dynamics, it is probably an accurate depiction as the absence of NMR observables is a consequence of conformational exchange between multiple conformations in solution. In the crystal, a well-defined conformation for loop 3 was observed, stabilised by extensive intermolecular crystal contacts. This conformation likely corresponds to one of the possible conformations adopted in solution.

**Figure 3.**
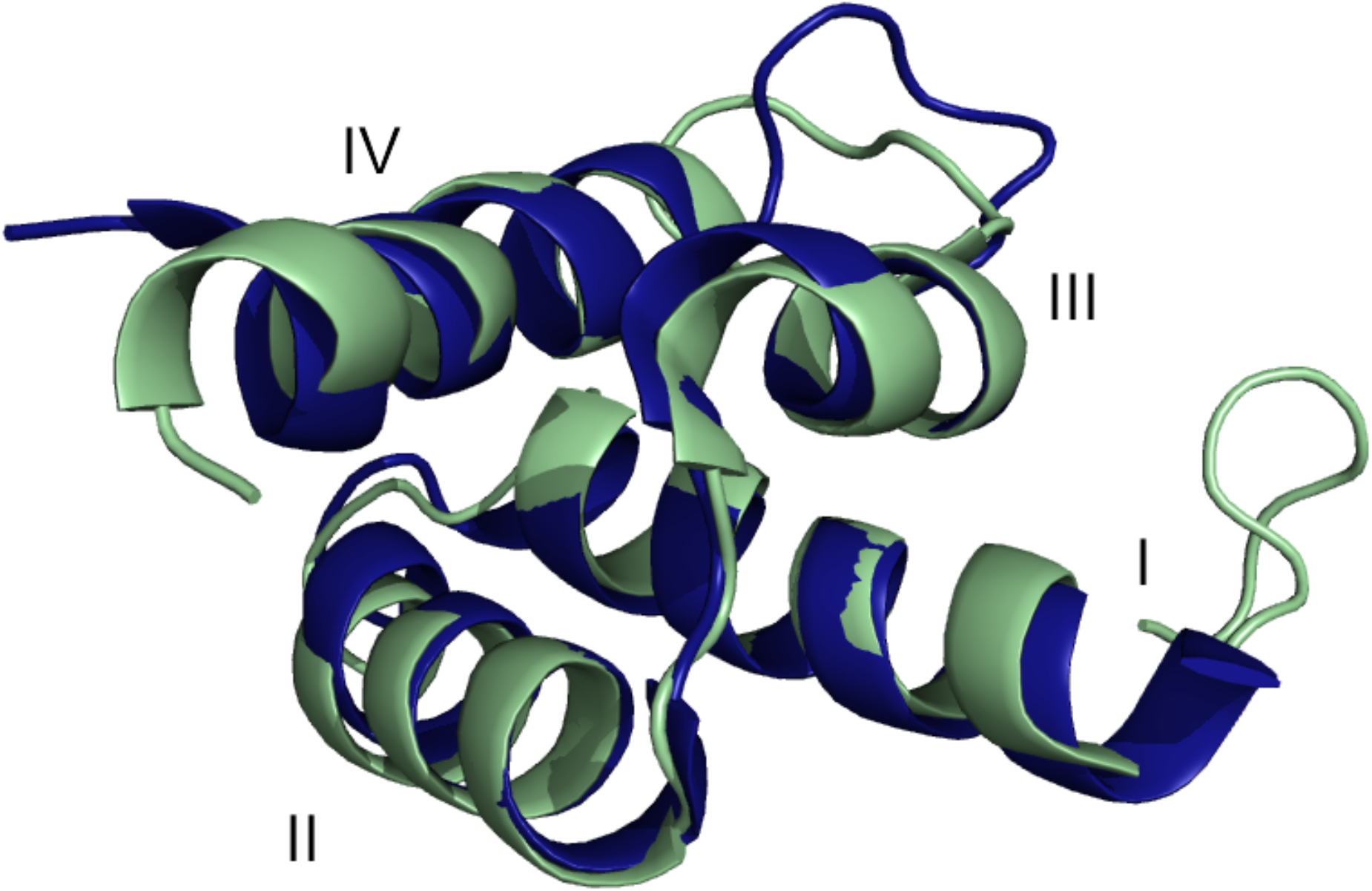
Structural comparison between the most representative AVR3a11_63-132_ structural model (model 5, pale green) and the AVR3a11_70-132_ crystal structure (blue, PBD 3ZR8 (9)). Helices are labelled with roman numerals. The RMSD between C_α_s in ordered regions of two structures is 1.05 Å. The main difference between the two structures is in the last C-terminal residues, due to intermolecular crystal contacts involving Tyr131.

The final three C-terminal residues of AVR3a11 emerge from helix 4 in the crystal structure and point away from the main body of the protein, stabilised by crystal contacts to Tyr131 (9). In solution, helix 4 extends further, bringing Tyr131 closer to helix 2. This conformation is more similar to the solution structure of another AVR3a homologue, *P. capsici* AVR3a4 (25). Outside the flexible regions (N-terminus and loop 3), there is also very good agreement between the solution structures of AVR3a11 and AVR3a4 (RMSD of 1.05 Å between Cα of the best representative models of each structural ensemble). Since the first 7 residues of AVR3a11_63-132_ were not well defined in the NMR structural models, studies of dynamics were performed on the AVR3a11_70-132_ construct.

### Hydrogen/deuterium exchange

To evaluate slow conformational dynamics in the structured AVR3a11_70-132_ effector domain, hydrogen/deuterium (H/D) exchange experiments were performed. H/D amide exchange rates are affected by the presence and stability of hydrogen bonds, and are influenced by protein fold stability and breathing (local unfolding) events. There was large variability in the exchange time for different residues. For 26 out of 63 residues, exchange was fast and completed in less than 4 minutes, before any NMR data could be acquired. On the other hand, Leu110, the residue with the slowest exchange, still retained a clear signal after 24 hours. For the residues where a peak was observed, protection factors were calculated (Fig. 4A). Protection factors, P = k_rc_/k_prot_, compare the exchange rate measured (k_prot_) with the exchange rate expected in a random coil with the same amino acid sequence (k_rc_). For residues with fast signal decay, protection factors could not be calculated, and were estimated as < 100, based on the average peak intensity before exchange and the signal-to-noise ratio of the spectra. A few residues could not be assigned, or have their exchange measured, due to peak overlap. Figure 4B illustrates the distribution of exchange times across the AVR3a11_70-132_ structure: fast exchanging residues are located mainly in helices 1 and 4, while slowly exchanging residues are located in helices 2 and 3, facing the hydrophobic core.

**Figure 4.**
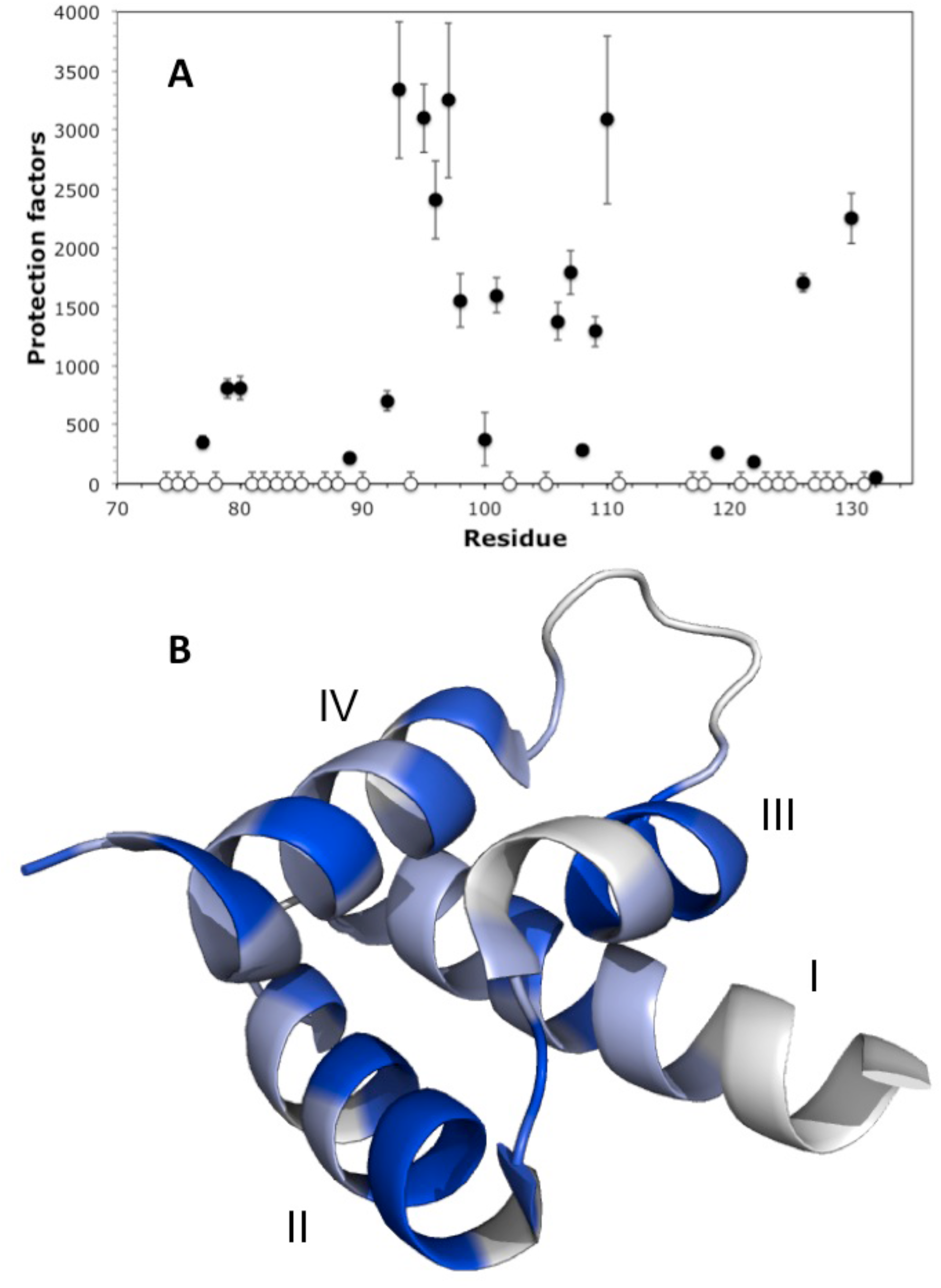
Hydrogen/deuterium exchange measurements for AVR3a11_70-132_ carried out at 5 °C and pH 6.8. A) Protection factors by residue. B) Crystal structure of AVR3a11_70-132_ coloured according to protection factor: larger than 1000 (dark blue), between 100 and 1000 (lighter blue), or too small to be measured (estimated under 100, lightest blue). Some residues could not be assigned (white). Helices are labelled with roman numerals.

The residues with the longest exchange times are generally associated with the structural core of the protein, in helices 2, 3 and 4. Residues in helix 1, which does not belong to the core WY folding unit (9), exchanged entirely in less than an hour and had protection factors lower than 1000. Of the previously described functionally relevant residues, the amides of Gln94 and Tyr131 both exchanged too fast for accurate protection factors to be calculated, while Glu71 could not be observed in any spectra, likely due to conformational broadening. Overall, the H/D exchange results for AVR3a11_70-132_ show a fairly stable structural core, corresponding to the WY-domain fold, surrounded by more dynamic regions, where the functionally relevant residues are located.

### Relaxation analysis

Fast (ps-ns) dynamics of the core region AVR3a11_70-132_ were analysed through backbone amide ^15^N NMR relaxation parameters at 500 and 800 MHz (Fig. 5). While the presence of conformational exchange clearly shows regions with enhanced dynamics in the ms-s timescale, and H/D exchange experiments reveal a stable structural core surrounded by more dynamic regions, in the ps-ns timescale AVR3a11_70-132_ shows uniform behaviour along the sequence, with the exception of the last C-terminal residue, Val132, which is significantly more flexible. However, residues near the exchange broadened loop 3, such as Lys111, have shorter T_1_, longer T_2_ and lower NOE values than other residues in the same region, suggesting greater flexibility in the proximity of the loop. A few residues could not be analysed, either because they were not assigned due to conformational broadening, or because they were overlapped in the [^1^H, ^15^N]-HSQC NMR spectra. Excluding Val132, average relaxation parameters for AVR3a11_70-132_ are T_1_ relaxation times of 456 ± 18 ms at 500 MHz and 733 ± 34 ms at 800 MHz; T_2_ relaxation times of 105 ± 8 ms at 500 MHz and 83 ± 8 ms at 800 MHz, and heteronuclear NOE ratios of 0.739 ± 0.046 at 500 MHz and 0.823 ± 0.044 at 800 MHz. T_1_/T_2_ ratios, often used to estimate an overall correlation time (τ_m_), were 4.36 ± 0.35 at 500 MHz, and 8.92 ± 0.97 at 800 MHz, corresponding to a τ_m_ of 6.9 ± 0.4 ns from the 500 MHz data, and 6.7 ± 0.4 ns from the 800 MHz data (62). Although the values at the two fields have very good agreement, they are significantly larger than the 5.8 ns that would be expected for an isotropic protein this size (63). Possible causes for this difference include anisotropy of the motions in the protein, some level of aggregation at the concentrations required by NMR, or more complicated motions.

**Figure 5.**
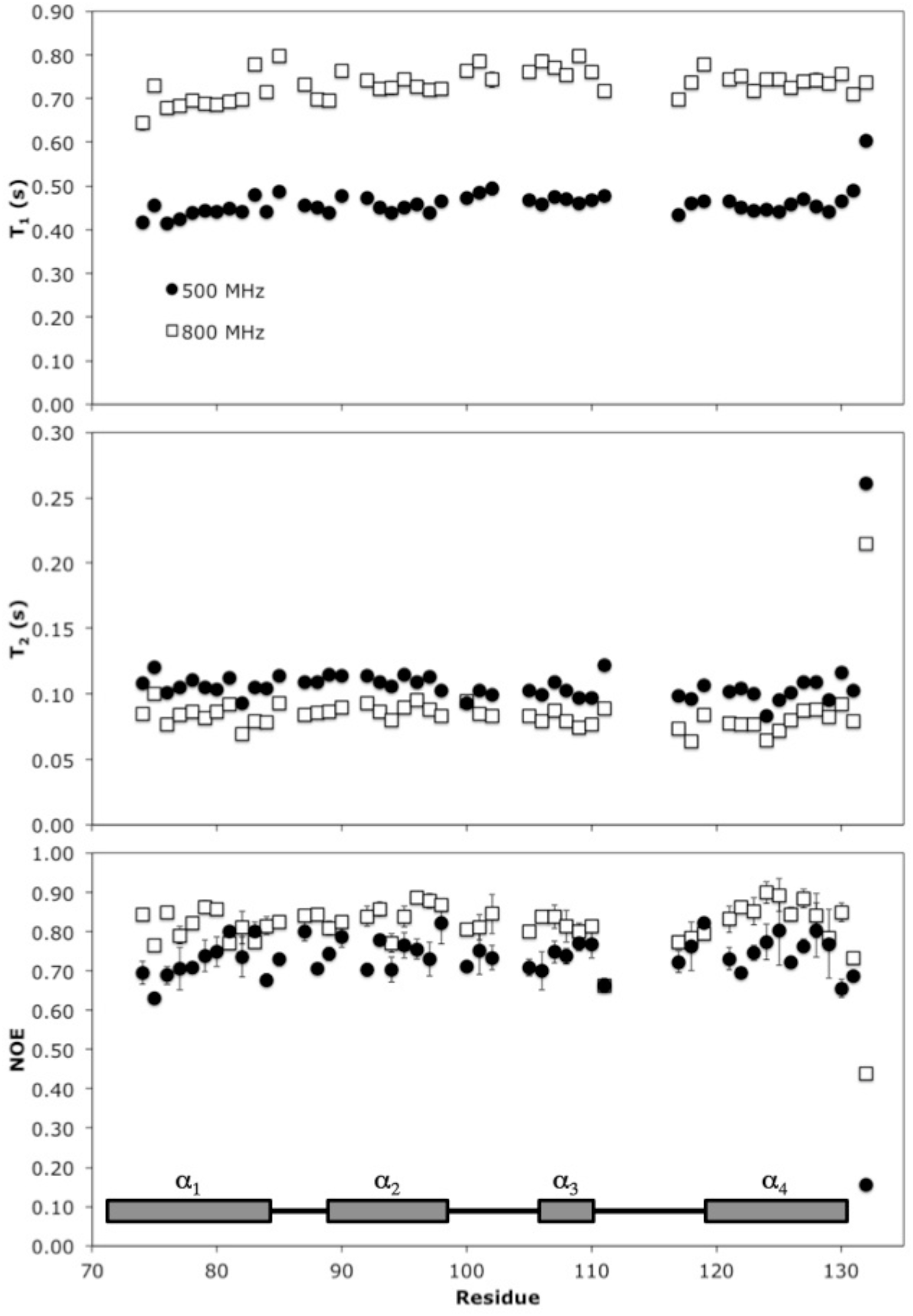
^15^N NMR relaxation parameters for AVR3a11_70-132_ plotted against residue number. T_1_ (top panel), T_2_ (middle panel) and heteronuclear NOE (bottom panel) were measured at 500 MHz (black circles) and 800 MHz (white squares). Secondary structure features are schematically represented at the bottom. Residues for which no data is available correspond to overlapped peaks or those which could not be assigned.

The data was fit to the well-established Lipari-Szabo model free approach (55, 56), showing extensive conformational exchange contributions throughout the protein, as indicated by the presence of R_ex_ terms for all residues fitted, with the exception of Val132 (Fig. S3). The presence of extensive conformational exchange undermines assumptions made during the model free fitting process, and therefore the values obtained are not reliable. For this reason, the data were further analysed with the reduced spectral density approach.

Relaxation parameters give information about motions in the protein through their relationship with the spectral density function, which describes the range and amplitude of frequencies sampled by each amide bond vector as the molecule reorients itself in the magnetic field. For rigid isotropic motion, the spectral density function J(ω) is given by equation 1:

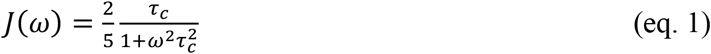

where ω is the frequency of motion and τ_c_ is the correlation time (64). For flexible molecules, J(ω) is a composite function, and can be expressed as the weighted sum (with appropriate scaling factors) of the spectral density functions of individual independent motions, with individual correlation times (65). For anisotropic molecules, the spectral density function for each amide bond vector will also be affected by the anisotropic tumbling of the molecule.

The reduced spectral density analysis approach (60) estimates the spectral density function, J(ω), at three different frequencies for each magnetic field used: 0, ω_N_ and 0.87ω_H_, corresponding to the contribution to the relaxation parameters of slow (ms-ns), intermediate (ns) and fast (ns-ps) motions, respectively, and where ω_N_ and ωH are the Larmor frequencies for ^15^N and ^1^H at that magnetic field. This approach was used to analyse the dynamics of AVR3a11_70-132_ (Fig. S4). J(0) is independently calculated from relaxation data acquired at each magnetic field, and therefore can be used to check the consistency of the data sets (66). The two data sets, at 500 and 800 MHz, have J(0) values within 2.5% of each other (Fig. S5), showing good consistency.

The estimated value of J(ω) throughout the protein sequence was fairly constant for J(ω_N_) at both magnetic fields, but showed greater variation for J(0) and J(0.87ωH), suggesting that different residues have variations in both fast internal motions (ps-ns timescale) and slow conformational exchange (ms-ns timescale) (Fig. S4). Graphical analysis can be used to interpret reduced spectral density mapping and relate it to the motions present in the protein (65, 67). The plot of J(0.87ω_H_) against J(ω_N_) (Fig. 6, top panel) is independent of slow conformational exchange. The solid lines represent the theoretical boundaries for the Lorentzian spectral density functions of completely rigid molecules with different overall correlation times, τ. Different regions of the plot correspond to different motional regimes, and are labelled A, B and C (68). Region A corresponds to τ under 300 ps, and residues in a protein found in this region would be dominated by fast motions. Region B corresponds to τ between 300 ps and 3 ns; and region C corresponds to τ longer than 3 ns. The data from AVR3a11_70-132_, in both fields, clusters close to the border of region C, suggesting that AVR3a11_70-132_ is a fairly rigid protein. At 800 MHz, the data points are clustered around a correlation time of about 6.8 ns, consistent with the values calculated from T_1_/T_2_ ratios. At 500 MHz, the data points correspond to a correlation time of approximately 7.8 ns, significantly larger than calculated from T_1_/T_2_ ratios. The isolated point towards the middle of the graphs corresponds to Val132, the more flexible last residue in the protein, whose dynamics can only be described by a combination of independent motions in different timescales.

**Figure 6.**
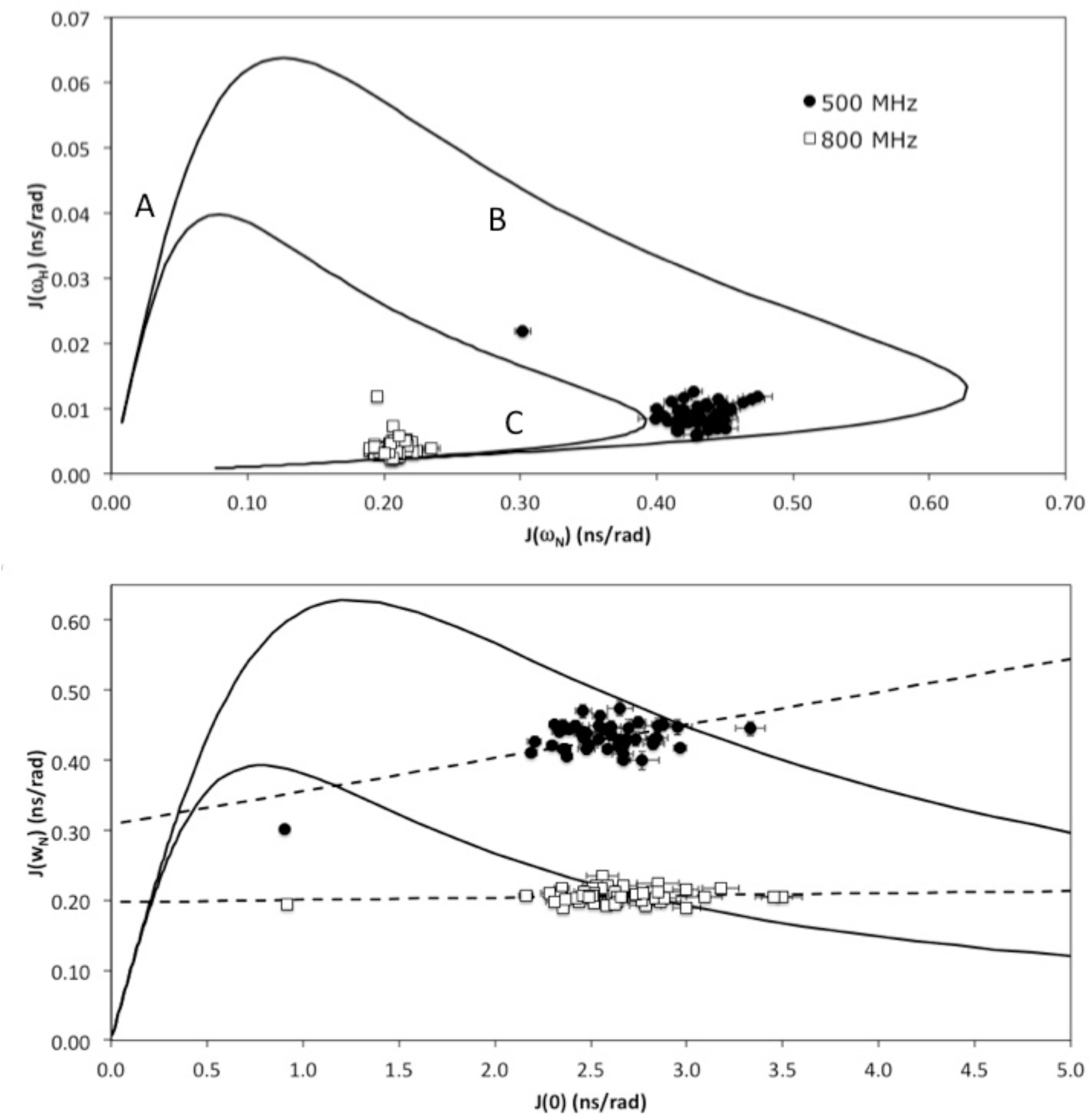
Plots for the dependency of J(ω_N_) on J(ω_H_) (top panel) and of J(ω_N_) on J(0) (bottom panel) at 500 MHz (black circles) and 800 MHz (white squares). The continuous lines were calculated for a simple Lorentzian spectral density function for a rigid particle at the corresponding magnetic fields. Dashed lines correspond to the least-squared fit for the data at each magnetic field. Data points outside the area delimited by the theoretical curve indicate the presence of slow motions characteristic of conformational exchange. Regions A, B, and C in the top panel correspond to different motional regimes.

The plot of J(ω_N_) against J(0) (Fig. 6, bottom panel) shows a linear relationship between J(ω_N_) and J(0), as expected from the theory. Again, the solid line represents a boundary of the combination of values that are theoretically possible for a rigidly tumbling molecule. The two intercepts between the theoretical curve and the line of best fit for each field correspond to values of τ of 0.6 ns and 6.8 ns, at 800 MHz, and 0.9 ns and 7.5 ns, at 500 MHz. For points located along each straight line, the larger value can be interpreted as the overall tumbling time, and the small value as corresponding to internal motions. Again, the correlation time calculated at 800 MHz is consistent with the values from T_1_/T_2_ ratios, while at 500 MHz the value is larger than from T_1_/T_2_ ratios. Additionally, we can observe that one of the values at 500 MHz, and multiple values at 800 MHz, are located outside the theoretical boundary. This suggests that AVR3a11_70-132_ does not conform to this simple model, and requires an extra term to account for slow motions, or conformational exchange. The greater number of residues that fall outside the theoretical curve at 800 MHz can be explained by the dependence of the conformational exchange term, R_ex_, on the magnetic field (69).

The graphical analysis of the reduced spectral density mapping shows that AVR3a11_70-132_ is dominated by motions somewhat slower than the overall tumbling time of the molecule, that affect an extensive region of the protein. This does not preclude the presence of fast motions, but it demonstrates that AVR3a11_70-132_ motions cannot be described by simple models that assume one overall tumbling time for the molecule combined with much faster internal motions.

### Conformational exchange

The multidimensional NMR spectra of both AVR3a11 constructs display signs of conformational exchange (Fig. 1 and S2), which are confirmed by the extensive presence of R_ex_ terms when the relaxation data is fitted to the Lipari-Szabo model-free approach. The effects of multiple exchanging protein conformations on NMR spectra depend on the rate of conformational exchange. When the exchange between conformations is slow, a peak will be observed for each conformation, with each peak intensity proportional to the corresponding population. When the rate of exchange (in s^-1^) is similar to the frequency difference between the corresponding NMR peaks (in Hz) this leads to broadening of the peaks, often beyond detection. If there is fast exchange between conformations, only one peak will be observed (corresponding to the weighted average of the contributions of each conformation), and the presence of conformational exchange may not be noticed. We observe both slow and intermediate exchange in the NMR spectra of AVR3a11.

Conformational exchange in the intermediate timescale is responsible for the broadening beyond detection of amide peaks for 15 amino acid residues, which could not be observed in the [^1^H, ^15^N]-HSQC NMR spectrum of AVR3a11_63-132_. These residues are located mainly in the N-terminal region of the construct (Gly63, Leu64, Asp66, Phe68-Glu71) and in loop 3 (Ser112-Gly116), with a few exceptions (Leu103, Thr104, Arg120).

Slow conformational exchange results in the presence of minor peaks both in the [^1^H, ^15^N]-HSQC and triple resonance spectra (Fig. S2) for residues Lys72, Lys76-Lys80, Gln94, Ser107, Lys111, Tyr125, Tyr128 and Val132. Residues Lys76 to Lys80 (in helix 1) are in close proximity to loop 3, and may reflect multiple conformations adopted by loop 3, rather than conformational variability in helix 1. Overall, most residues affected by conformational exchange are outside the core WY-domain fold (Fig. S6).

The separation between major and minor conformation peaks in AVR3a11_63-132_ ranges from 0.043 to 0.267 ppm in the ^1^H dimension. In the spectrometer used in these experiments, these correspond to frequencies between 34.41 and 213.66 Hz. For peaks to be clearly observed in slow exchange, it can be seen from simulations that the exchange rate should be at least 10 times slower than the difference in frequency (70), and therefore in AVR3a11_63-132_ the backbone conformational exchange rates for slowly exchanging residues are slower than ≈ 3.5 s^-1^, which is consistent with the largest R_ex_ values fitted in the model-free approach.

## Discussion

To date, no function has been ascribed to *P. capsici* AVR3a11. However, its crystal and solution structures contribute to the general understanding of WY domain effectors, which include *P. infestans* AVR3a and other highly similar proteins across oomycete species. The key allelic variant positions in AVR3a, residues 80 (Glu/Lys) and 103 (Ile/Met), which are involved both in recognition by R3a and PCD suppression, correspond to AVR3a11 Glu71 and Gln94. Gln94 is positioned in the middle of helix 2 (Fig. 2), and could not be completely assigned in the NMR spectra due to conformational exchange. This highlights the conformational variability of these regions, and suggests that functionally relevant residues may be located in dynamic regions. Very short T_2_ relaxation times were observed for Glu80 in Avr3a_48-147_ (14), suggesting that conformational exchange for these residues is conserved across homologous proteins.

AVR3a homologues show greater sequence variation in the C-terminal end. In AVR3a, this region contains Tyr147, which has a role in PCD suppression but not R3a recognition. While it could be expected that the tyrosine near the C-terminus has a similar role in effectors with the same fold, in the solution structure of *P. capsici* AVR3a_60-147_ (14) Y147 seems to be in a flexible region, and not at the end of a helix, as seen for tyrosines in AVR3a11 and AVR3a4. However, the presence of extra residues beyond the end of the AVR3a sequence in the construct used make it difficult to judge the natural conformation for helix 4 in AVR3a.

In addition to the crystal structure of AVR3a11 and the solution structures of AVR3a and AVR3a4, other structures of WY-domain containing effectors from oomycetes have been previously described, such as PexRD2 from *P. infestans* (9) and ATR1 from *Hyaloperonospora arabidopsidis* (71), which contains three WY domains. Comparing those structures, loop 3 shows the greatest sequence and structural diversity, varying from 7 to 24 residues, and from unstructured to helical. In the solution structure of AVR3a11_63-172_, the disordered loop confirms that loop 3 corresponds to a region capable of adopting different conformations in WY-domains. Loop 3 is also missing assignments for a couple of residues in AVR3a_60-147_ (14) (BMRB accession code 25944) and ^15^N assignments for a couple of residues in AVR3a4 (25) (BMRB accession code 17588). Residues N-terminal to the effector domain and within loop 3 in AVR3a4 were reported to show conformational flexibility (25). This type of broadening was also observed for an internal disordered loop in the *H. arabidopsidis* RXLR effector ATR13 (which does not appear to contain a WY domain) (72).

The ability to sample multiple conformations may be relevant for the functional role of some proteins: in ATR13, the broadened loop is involved in localisation to the nucleolus (72), and in *P. infestans* AVR3a, a number of gain-of-function mutations that allow the activation of R3a HR by the AVR3a^EM^ isoform (17) were mapped to loop 3 (25).

Using H/D exchange and NMR relaxation analysis, we observed dynamics in the effector domain of AVR3a11 at different timescales. Motions in the ms range and slower dominate the dynamics of AVR3a11_70-132_, as seen from graphical analysis of the reduced spectral density mapping. Hydrogen/deuterium exchange data indicate a stable hydrophobic core which excludes helix 1 and loop 3, with residues Tyr93, Lys95-Asp98 (helix 2), Tyr101 (loop2), Leu106, Ser107, Phe109, Leu110 (helix 3), Met126 and Asp130 (helix 4) showing high protection factors. These residues include most of the stabilising interactions described for the crystal structure (9). The first few residues in helix 4 (including the conserved Tyr125) are less rigid than expected, possibly affected by loop 3. This view is consistent with the signs of conformational exchange observed, in which residues in helix 1 and N-terminal to it, jointly with loop 3 and a few other residues, were strongly affected by slow and intermediate chemical exchange.

H/D exchange rates for residues in slow conformational exchange (showing the presence of multiple HN crosspeaks) were variable, with protection factors ranging from 1600 to too small to be measured. The lack of correlation between conformational exchange measured by H/D exchange and by the presence of multiple crosspeaks is not surprising, given the different timescales of conformational changes measured by each technique. Multiple conformations in exchange at a rate of 3.5 s^-1^ should give rise to multiple peaks, while still more than a thousand times faster than the limit of detection for our H/D exchange experiments.

While our combined experiments allow us to determine the presence of slow (conformational exchange) motions, they do not provide any information as to their nature. Limited relaxation experiments with varying concentrations (2-fold) and magnetic fields suggested a dependence on field intensity and not on concentration (data not shown). This makes it very unlikely that transient intermolecular interactions are involved. Additionally, we have previously shown AVR3a11 to be a monomer in solution, with no evidence of partial dimerization in either gel filtration or analytical ultracentrifugation experiments (9). Therefore, the general presence of conformational exchange in AVR3a11_70-132_ suggests that those intramolecular motions could correspond to large conformational changes or partial unfolding of the protein.

In summary, the effector domain of AVR3a11 is a small four-helix bundle in solution, with a stable hydrophobic core, which is preserved in the solution structure despite the highly dynamic characteristics of this protein and the addition of 7 N-terminal residues to AVR3a11_70-132_. The dynamics of AVR3a11_70-132_ are dominated by slow motions, as evident from NMR relaxation measurements, from the presence of peaks corresponding to minor conformations in the NMR spectrum, and from NMR peaks broadened beyond detection. As functionally important residues are found in regions with extensive conformational exchange, and conformational exchange was observed in other WY-domain effectors, flexibility may have a functional role in this family of effectors.

H/D exchange results reveal that the structures of helices 2, 3 and 4 are more stable than that of helix 1. This reinforces the idea that the folding core is formed by the 3-helical WY-bundle, a widespread structural unit in RXLR effectors (24, 73), with the addition of helix 1 as one of several possible adaptations for stability and function (9). We predict that this increased dynamic stability for the three core helices in the WY-domain is also likely to be encountered in other effectors of this family.

## Abbreviations

AVR: avirulence protein
H/D: hydrogen/deuterium
HR: hypersensitive response
NLR: nucleotide binding-leucine rich repeat
NMR: nuclear magnetic resonance
NOE: nuclear Overhauser effect
PAMP: pathogen-associated molecular pattern
PCD: programmed cell death
PDB: Protein Data Bank
PRR: pattern recognition receptor
R3a: resistance protein 3a

## Author contributions

JT, VSC and LSB performed the research and analysed the data. MJB designed the research and contributed experimental tools. TMAB designed the research, analysed the data and wrote the manuscript, with input from the other authors.

## Acknowledgements

We thank Dr Colin MacDonald for his support in maintaining the NMR spectrometers, and Dr Mark Howard (currently at Bruker) and Prof Geoff Moore (University of East Anglia) for helpful discussions.

JT’s PhD studentship was funded by the Biotechnology and Biological Sciences Research Council (UK).

