## Supplemental Material for "Dynamics in the *Phytophthora capsici* effector AVR3a11 confirm the core WY domain fold"

Supplementary material

**Figures**

Figure S1. AVR3a11$_{63-132}$ sequence coloured by assignment status: fully assigned (dark green), side chain assigned but not backbone atoms (light green), partially assigned (orange) and not visible in the spectra (grey).

GLTDLFKTEK AAVKKMAKAI

MADPSKADDV YQKWADKGYT

LTQLSDFLKS KTRGKYDRVY

NGYMTYRDYV

Figure S2. Detail of regions in the [$^1$H, $^{15}$N]-HSQC two-dimensional NMR spectrum of AVR3a11$_{63\text{-}132}$, showing small peaks corresponding to minor conformations of AVR3a11$_{63\text{-}132}$ for Lys111 (top panel), Lys 80 (middle panel), Lys72 and Lys 76 (bottom panel). All peaks were assigned with the aid of three-dimensional triple resonance spectra.

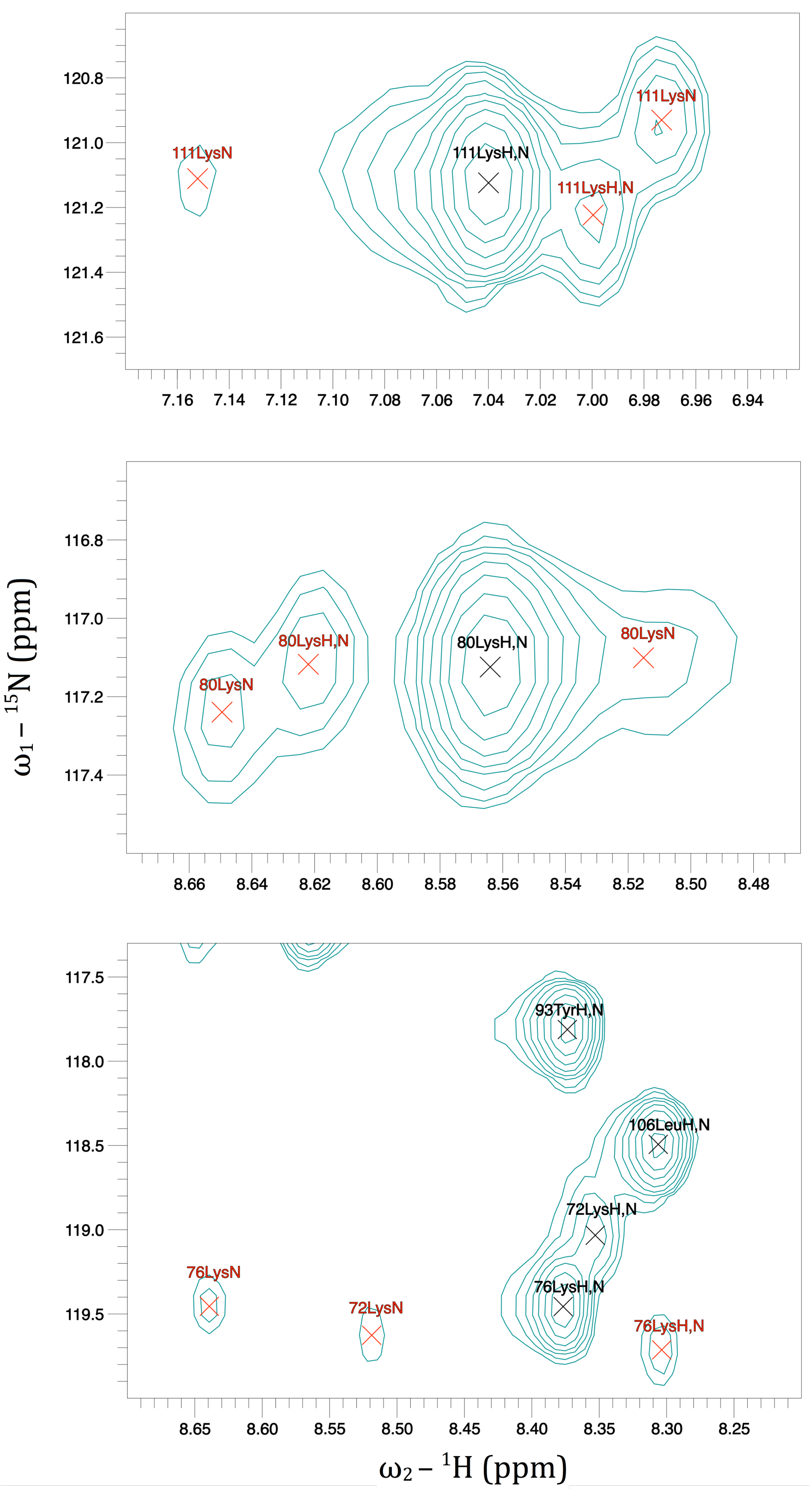

Figure S3. Model-free analysis of AVR3a11$_{70\text{-}132}$ relaxation plotted against residue number, displaying the order parameter $S^2$ (top), effective correlation time for internal motions $\tau_e$ (middle), and exchange rate $R_{ex}$ (bottom). Data points are coloured by the model used to fit each residue, in which M2 (white) uses $S^2$ and $\tau_e$, M3 (black) uses $S^2$ and $R_{ex}$, M4 (grey) uses $S^2$, $\tau_e$ and $R_{ex}$, and M5 (striped) uses $S_s^2$, $S_f^2$, $\tau_e$ and $R_{ex}$, where $S_s^2$ and $S_f^2$ are, respectively, slow and fast order parameters. Two residues (Ala79 and Leu106) fitted as M5 yielded very large, inaccurate $\tau_e$ values that were excluded from the plot. Error bars represent the fit error, and are too small to be seen in the order parameter plot.

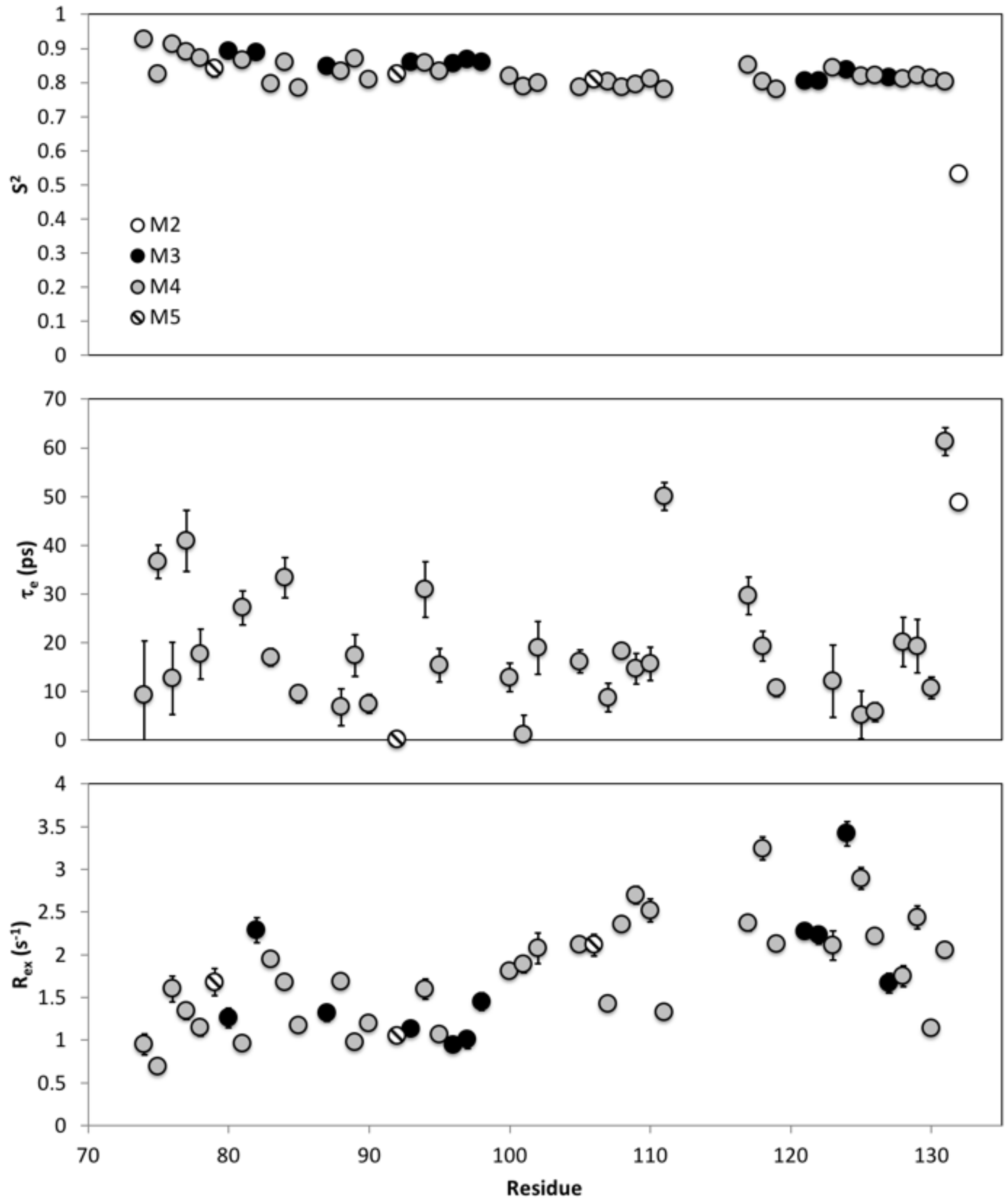

Figure S4. Reduced spectral density analysis for AVR3a11$_{70-132}$ plotted against residue number. Frequencies analysed correspond to slow [J(0), top panel], intermediate [J($\omega_N$), middle panel] and fast [J($\omega_H$), bottom panel] motions, calculated from data at 500 MHz (black circles) and 800 MHz (white squares).

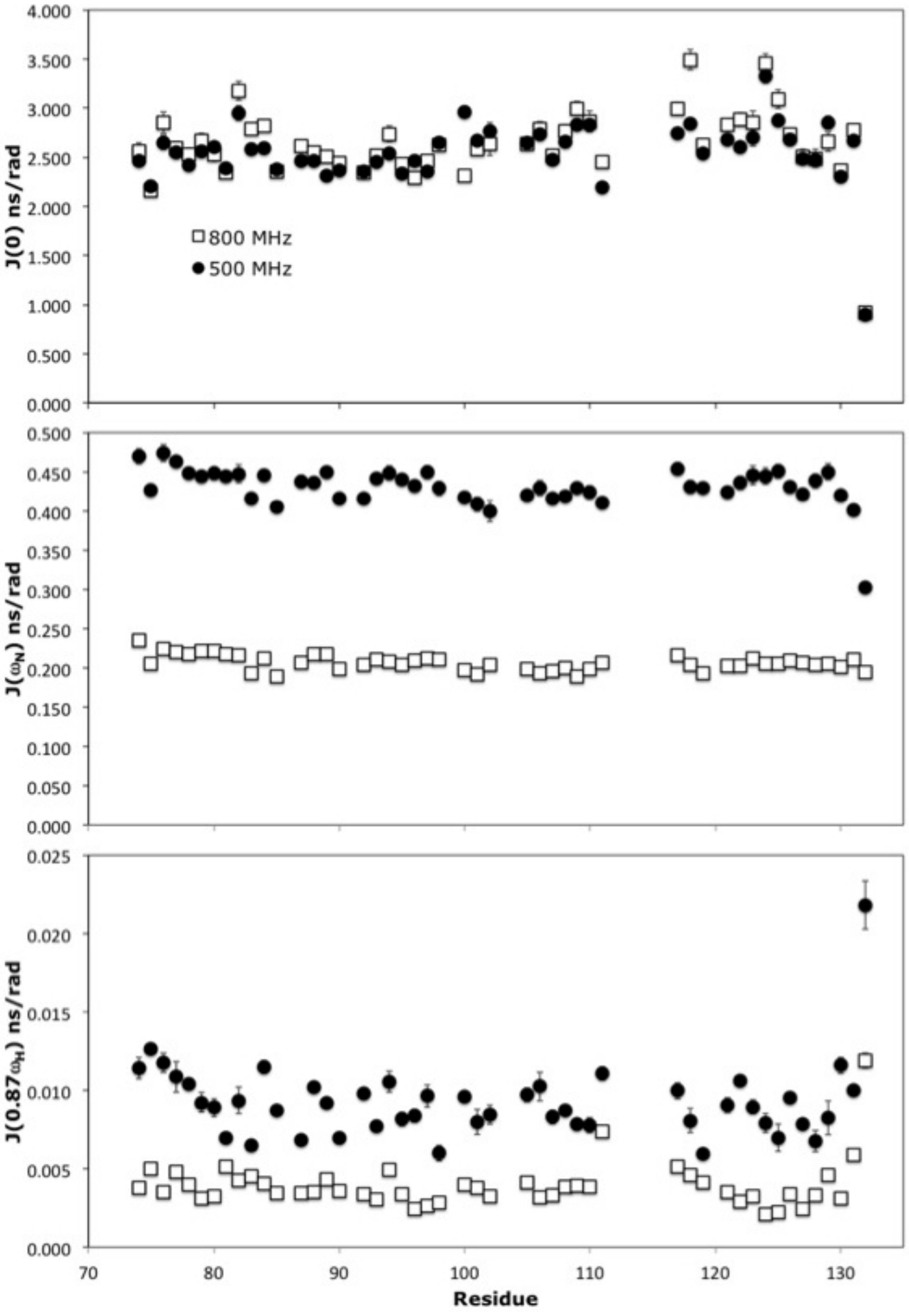

Figure S5. Correlation between J(0) calculated from data at 500 MHz and 800 MHz, to confirm consistency between the two data sets. Residues 100 and 118 show the largest differences in J(0) values.

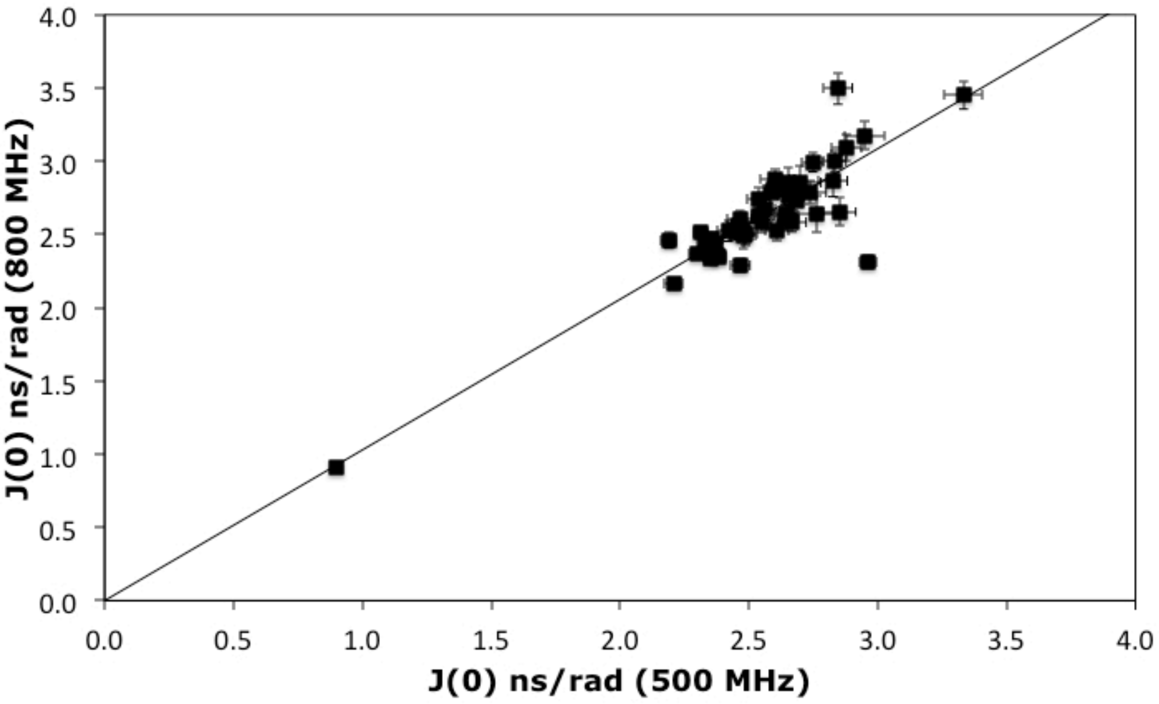

Figure S6. Most representative conformer of AVR3a11$_{63\text{-}132}$ solution structure, coloured according to evidence for conformational exchange: residues with peaks broadened beyond detection in the [$^{1}$H, $^{15}$N]-HSQC NMR spectrum (grey) and residues with minor peaks in the [$^{1}$H, $^{15}$N]-HSQC NMR spectrum, corresponding to alternative conformations (blue).

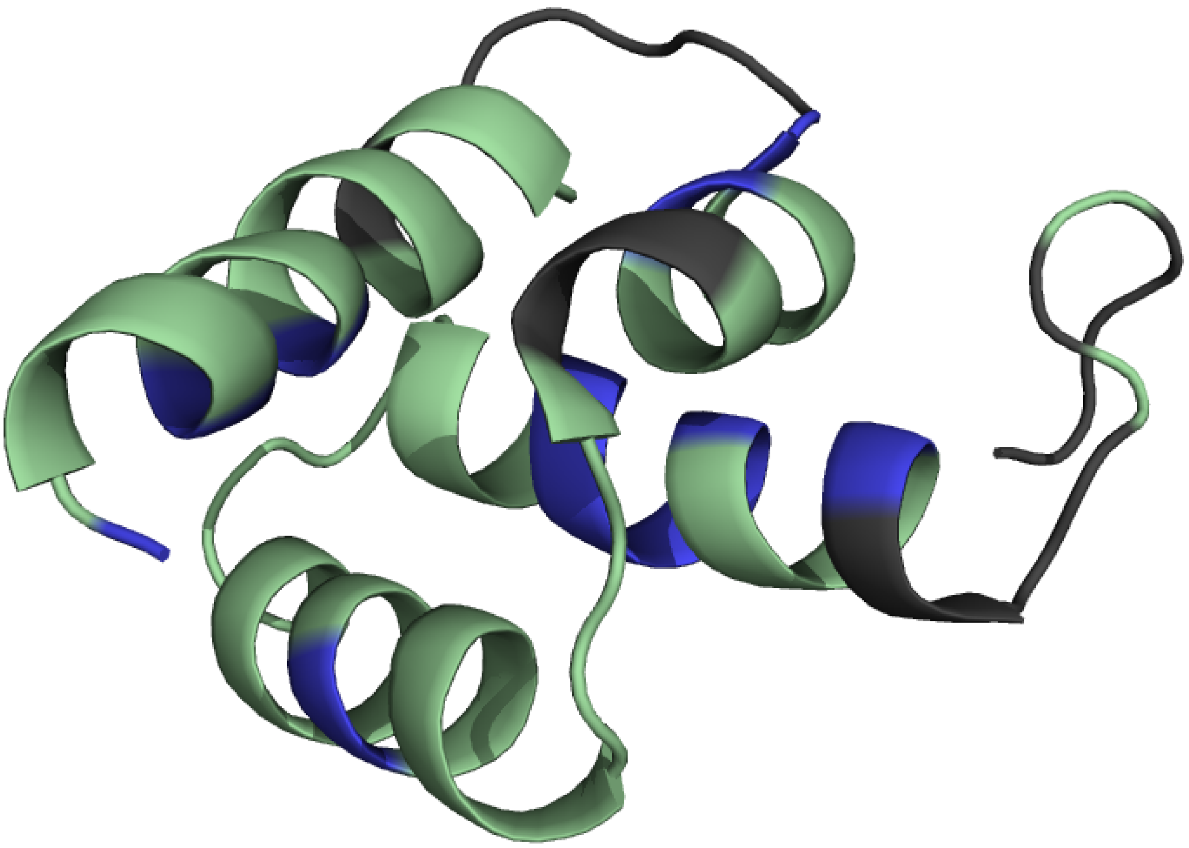

Table S1. Experimental restraints and structural statistics for AVR3a11$_{63-132}$.

| | |
|---|---|
| **Distance restraints** | |
| All | 870 |
| Intra-residue | 158 |
| Sequential | 227 |
| Medium range ($1 < \|i - j\| \leq 4$) | 296 |
| Long range ($\|i - j\| > 4$) | 195 |
| **Dihedral angle restraints** | 98 |
| **Residual constraint violations** | |
| NOE violations > 0.2 Å | 0 |
| Dihedral angle violations > 5° | 0 |
| Van der Waals violations > 0.1 Å | 0 |
| **Backbone deviation from average structure (RMSD)**[a] | |
| All residues | 2.9 ± 0.7 Å |
| Ordered | 0.75 ± 0.12 Å |
| **Ramachandran plot**[a,b] | |
| Most favoured regions | 94.40% |
| Additionally allowed regions | 5.60% |
| Generously allowed or disallowed regions | 0.00% |

[a]Calculated using the validation software PSVS [47].
[b]Ordered residues, calculated using Procheck [49].
